# Input-specific NMDAR-dependent potentiation of dendritic GABAergic inhibition

**DOI:** 10.1101/155077

**Authors:** Chiayu Q. Chiu, James S. Martenson, Maya Yamazaki, Rie Natsume, Kenji Sakimura, Susumu Tomita, Steven J. Tavalin, Michael J. Higley

## Abstract

Preservation of a balance between synaptic excitation and inhibition is critical for normal brain function. A number of homeostatic cellular mechanisms have been suggested to play a role in maintaining this balance, including long-term plasticity of GABAergic inhibitory synapses. Many previous studies have demonstrated a coupling of postsynaptic spiking with modification of perisomatic inhibition. Here, we demonstrate that activation of NMDA-type glutamate receptors leads to input-specific long-term potentiation of dendritic inhibition mediated by somatostatin-expressing interneurons. This form of plasticity is expressed postsynaptically and requires both CaMKIIα and the β2-subunit of the GABA-A receptor. Importantly, this process may function to preserve dendritic inhibition, as *in vivo* loss of NMDAR signaling results in a selective weakening of dendritic inhibition. Overall, our results reveal a new mechanism for linking excitatory and inhibitory input in neuronal dendrites and provide novel insight into the homeostatic regulation of synaptic transmission in cortical circuits.

## Introduction

The balance of synaptic excitation and inhibition is central to normal brain function and is disrupted in a variety of neurodevelopmental disorders (Gogolla et al., 2009; Isaacson and Scanziani, 2011; Lewis and Hashimoto, 2007). In the neocortex, this balance is hypothesized to be maintained via an array of mechanisms that regulate synaptic strength and excitability (Kullmann et al., 2012; Malenka and Bear, 2004; Turrigiano, 2011). Mechanistic studies of synaptic plasticity have largely focused on potentiation and depression of excitatory glutamatergic connections. More recently, plasticity of inhibitory GABAergic synapses has also begun to receive attention, although the underlying cellular targets and molecular mechanisms are less well understood (Castillo et al., 2011; Kullmann et al., 2012).

A major challenge to understanding the contribution of inhibitory plasticity to brain development and function is the diversity of cortical GABAergic interneurons (INs)(Ascoli et al., 2008). Recent work suggests three principal groups: cells co-expressing either the calcium (Ca2+) binding protein parvalbumin (PV), the peptide transmitter somatostatin (SOM), or the serotonin 5HT3a receptor (Rudy et al., 2011). The latter class includes the vasoactive intestinal peptide (VIP)-expressing cells. PV-INs make inhibitory contacts onto the perisomatic and proximal dendritic regions of excitatory pyramidal neurons (PNs) and exert well-documented control over the magnitude and timing of PN spike output (Cardin et al., 2009; Pouille and Scanziani, 2001). SOM-INs contact dendritic arbors where they regulate Ca2+ signaling, synaptic integration, and dendritic spikes (Chiu et al., 2013; Murayama et al., 2009). VIP-INs largely, though not exclusively, target other INs and may drive state-dependent disinhibition of PNs (Fu et al., 2014; Pfeffer et al., 2013).

Recent evidence using 2-photon imaging of fluorescently tagged inhibitory synapses *in vivo* suggests distinct learning rules for different populations of GABAergic inputs (Chen et al., 2012; Villa et al., 2016). In particular, inhibitory synapses onto dendritic spines, potentially formed by SOM-INs (Chiu et al., 2013), appear to be particularly plastic, as their basal turnover and response to sensory deprivation is significantly more dynamic than those onto dendritic shafts (Chen et al., 2012; van Versendaal et al., 2012). These findings suggest the intriguing possibility of GABAergic circuit-specific plasticity.

Notably, most studies of GABAergic plasticity have implicated perisomatic inhibition as a key locus for regulation. For example, synapses formed by PV-INs in primary visual cortex selectively exhibit long-term potentiation (iLTP) in response to activity-dependent release of nitric oxide by postsynaptic PNs (Lourenco et al., 2014), and inputs from fast-spiking, putative PV-INs, onto layer 4 PNs are selectively modified by visual experience (Maffei et al., 2006). Similarly, cholecystokinin (CCK)-expressing basket cells targeting proximal somatodendritic regions in the hippocampus are particularly sensitive to retrograde endocannabinoid signaling (Lee et al., 2010). Finally, Purkinje cell-targeting basket cells in the cerebellum exhibit iLTP in response to postsynaptic Ca2+ signaling (He et al., 2015). It is less clear whether GABAergic inputs to neuronal dendrites are regulated by similar mechanisms. Because excitatory and inhibitory synapses are in close proximity within dendritic compartments, glutamatergic activity may intimately shape dendritic inhibition. Indeed, previous studies in cultured hippocampal neurons, where circuit architecture is not preserved, suggested links between Ca2+ influx through NMDA-type glutamate receptors (NMDARs) and GABAergic synaptic function (Marsden et al., 2007; Petrini et al., 2014).

To determine whether glutamatergic signaling can directly influence inhibitory synaptic potency in intact cortical circuits, we utilized optogenetic stimulation of targeted GABAergic IN populations paired with activation of postsynaptic NMDARs. Our results show the remarkable finding that Ca2+ influx through NMDARs selectively drives iLTP of SOM-IN synapses, while inputs from PV-INs and VIP-INs are unaffected. This form of plasticity is expressed postsynaptically and requires the β2 subunit of the GABA_A_ receptor, which functions preferentially at SOM-IN synapses. Finally, we show that disruption of NMDAR activity *in vivo* leads to distinct consequences for perisomatic and dendritic inhibition. These results demonstrate molecular heterogeneity of GABAergic synapses in different somatodendritic compartments that corresponds to the presynaptic partner. This heterogeneity has direct consequences for activity-dependent, homeostatic balancing of excitatory and inhibitory circuits.

## Results

To examine the impact of glutamatergic signaling on specific subsets of GABAergic synapses, we used a viral vector to conditionally express EYFP-fused channelrhodopsin-2 (ChR2) in three populations of GABAergic interneurons (SOM-, PV-, or VIP-INs) within the mouse medial prefrontal cortex (Figure 1A-C, left). We selectively activated ChR2-expressing cells in acute slices with brief pulses of 473-nm light while monitoring the corresponding inhibitory postsynaptic currents (IPSCs) in nearby L2/3 PNs (Figure 1A-C, middle). In these experiments, PNs were loaded with chloride through the patch pipette to obtain detectable inward IPSCs at a holding potential of −70 mV. After obtaining a stable baseline, 20 μM NMDA was bath applied for 2 minutes and rapidly washed out (Figure 1A-C, right). In all experiments, inhibitory currents disappeared during NMDA wash-in and reappeared in the first 2 minutes after NMDA cessation, presumably due to NMDA-induced depolarization block of presynaptic neurons. Experiments using SOM-INs revealed that chemical activation of NMDARs produced a significant potentiation of optically evoked IPSCs (SOM-IPSCs), reaching a plateau ^~^20 min after NMDA washout (171 ± 18%, n = 8 cells, p=0.02; Figure 1A, E). This rise was not correlated with changes in series or membrane resistance (Figure 1D). Surprisingly, inhibitory responses mediated by either PV- (PV-IPSCs) or VIP-interneurons (VIP-IPSCs) did not exhibit potentiation following NMDAR activation, only recovering back to baseline (PV: 105 ± 5%, n = 7 cells, p=0.78; VIP: 91 ± 8%, n = 8 cells, p=0.20). Thus, our results demonstrate that glutamatergic signaling can drive iLTP in the neocortex, but this phenomenon is specific to a subpopulation of GABAergic synapses.

**Figure 1.**
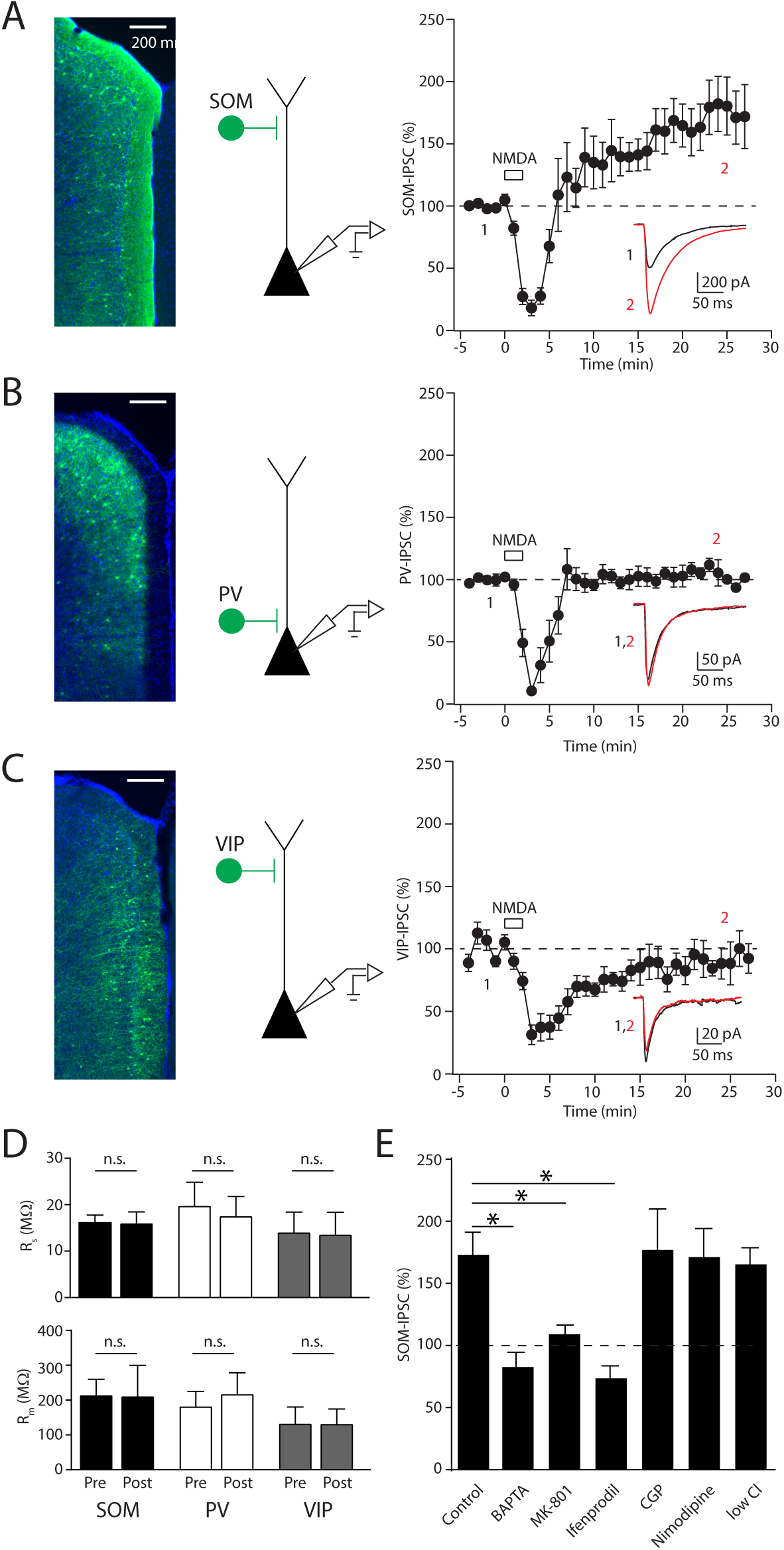
NMDA selectively potentiates GABAergic inhibition mediated by SOM-INs. (A-C, left) Epifluorescence images of ChR2-EYFP (green) expression and DAPI (blue) in the prefrontal cortex of the three different IN-Cre driver lines. (A-C, middle) Schematic of the recording and stimulation conditions. (A-C, right) Time-course of inhibitory postsynaptic currents in L2/3 pyramidal cells evoked by photo-activation of the specific interneuron types before and after brief application of 20mM NMDA. Average IPSC traces obtained before (black) and after (red) NMDA exposure from a single experiment at time points indicated are shown in the insets. (D) Series (top) and membrane resistance (bottom) are unchanged after brief NMDA exposure for all three groups. (E) Summary plot of the involvement of different Ca2+ sources and receptors on iLTP of SOM-IN inputs. Asterisks denote p-value of <0.05, Mann-Whitney Test.

Bath-application of NMDA may increase neuronal activity in the slice, leading to release of unspecified transmitters that might mediate iLTP. Therefore, we determined the requirement for postsynaptic NMDAR signaling in the recorded PN by loading cells with the NMDAR blocker MK-801 through the patch pipette. This manipulation abolished iLTP of SOM-IPSCs (109 ± 8%, n = 7 cells, p=0.005 compared to control; Figure 1E). In particular, NMDARs containing GluN2B-subunits are required for iLTP, as bath-application of the specific antagonist ifenprodil also blocked plasticity (74 ± 11%, n = 5 cells, p=0.02 compared to control; Figure 1E). Moreover, chelating postsynaptic Ca2+ by including BAPTA in the patch pipette also blocked iLTP (83 ± 10%, n = 4 cells, p=0.001 compared to control; Figure 1E). These results strongly indicate that iLTP is induced cell-autonomously by the activation of postsynaptic NMDARs and subsequent Ca2+ influx.

Additional pharmacological assays revealed that blockade of either GABA_B_ receptors (CGP-55845: 177 ± 33%, n = 3 cells, p=0.84) or L-type voltage-gated Ca2+ channels, (nimodipine: 171 ± 23%, n = 4 cells, p=0.60) did not reduce the magnitude of iLTP compared to controls. In addition, we confirmed that iLTP is also observed when monitoring outward currents at +10 mV in cells containing a physiological chloride concentration (low chloride: 165 ± 14%, n = 6 cells, p=0.89 compared to high chloride control), arguing that plasticity is not due to a change in the GABA_A_ reversal potential and is not an artifact of chloride-loading (Figure 1E).

NMDAR-dependent plasticity is often linked to Ca2+-dependent activation of CaMKIIα. Therefore, we tested the role of this kinase in iLTP. Both extracellular blockade with the antagonist KN-62 and intracellular blockade by cell-loading with autocamtide-2-related inhibitory peptide (AIP) abolished iLTP of SOM-IPSCs (control: 156 ± 16%, n = 8 cells; KN-62: 91 ± 5%, n = 7 cells, p=0.0003; AIP: 100 ± 5%, n = 8 cells, p=0.0002; Figure 2A, B).

**Figure 2.**
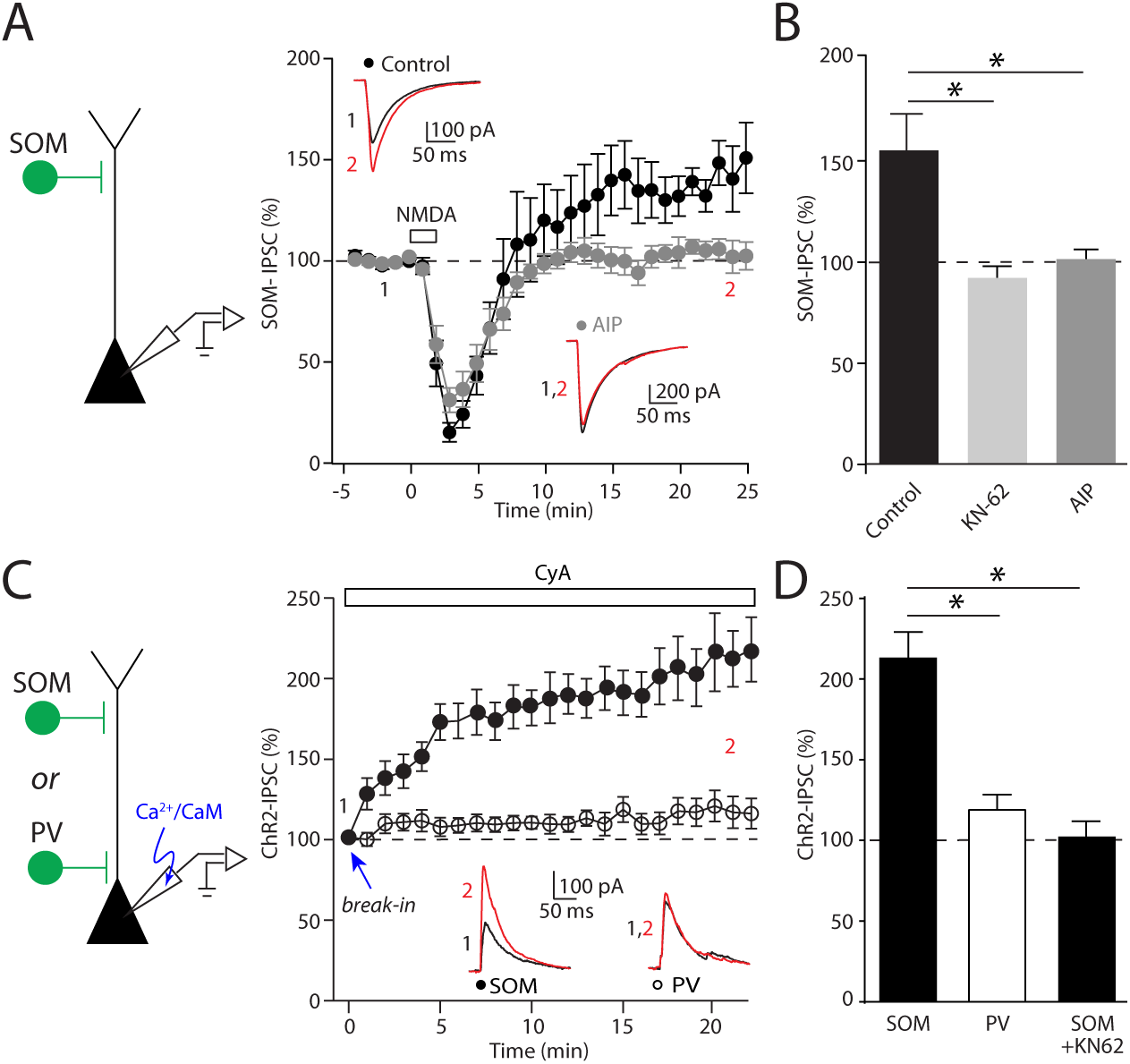
CaMKIIα activity mediates iLTP of dendritic GABAergic inhibition. (A) Time-course of inhibitory postsynaptic currents evoked by photo-activation of SOM-INs before and after NMDA application using control internal solution or patch solution loaded with AIP. Average IPSC traces obtained before (black) and after (red) NMDA exposure from a single experiment at time points indicated are shown in the insets. (B) Summary plot of the effect of blocking CaMKIIα on iLTP with bath applied KN-62 or cell loaded AIP. (C) Time-course of inhibitory postsynaptic currents evoked by photo-activation of SOM or PV-INs immediately after whole-cell break-in using an internal patch solution that contains calcium and calmodulin. To isolate the effect of CaMKIIα, the calcium-sensitive phosphatase, calcineurin, was blocked with bath application of cyclosporin A (CyA) throughout the experiment. Average IPSC traces obtained in the first minute (black) and after twenty minutes (red) of whole cell patch recording from a single experiment are shown in the insets. (D) Summary plot of the effect of loading calcium and calmodulin on the amplitude of inhibitory responses elicited by optogenetic activation of SOM- and PV-INs. The IPSC increase observed for SOM-IN inputs is abolished by KN-62. Asterisks denote p-value of <0.05, Mann-Whitney Test.

The lack of CaMKIIα-dependent iLTP at PV-IN synapses might reflect either absence of kinase at these perisomatic inputs or insensitivity to its actions. To distinguish between these possibilities, we examined whether direct activation of CaMKIIα is sufficient to potentiate IPSCs. In initial experiments, cells were loaded with Ca2+ and calmodulin through the patch pipette in the presence of the calcineurin antagonist cyclosporine A (Wang and Kelly, 1995). We began by recording IPSCs evoked by optical stimulation of SOM-INs immediately after breaking into the cell and observed a steady augmentation of response amplitude (213 ± 16%, n = 10, p=0.002), which was not observed in cells loaded with control pipette solution (122 ± 10%, n = 7, p=0.22; Figure 2C,D). KN-62 abolished the effect of loading Ca2+/calmodulin (103 ± 11%, n = 6, p=0.0005; Figure 2D), suggesting that direct activation of CaMKIIα is sufficient to potentiate inputs from SOM-INs. In striking contrast, loading the cell with Ca2+ and calmodulin had no effect on IPSCs evoked by stimulating PV-INs (Ca2+calmodulin: 120 ± 9%, n = 8, p=0.20; control: 112 ± 15%, n = 8, p=0.31, Fig. 2C, D). We confirmed the specificity of these findings by repeating similar experiments, but this time loading the PN with a constitutively active CaMKIIα (10 nM CaMKII*)(Tavalin and Colbran, 2017). Again, the amplitude of IPSCs mediated by SOM-INs but not PV-INs increased 20 min after whole-cell break-in (SOM: 146 ± 10%, n = 5 cells, p=0.02; PV: 108 ± 12%, n = 5 cells, p=0.58; Supplemental figure 1). Together, these results indicate that the inherent sensitivity to CaMKIIα signaling differs across distinct GABAergic synaptic populations.

Our results suggest that iLTP induction requires postsynaptic NMDARs, Ca2+ influx, and CaMKIIα activation. However, the site of expression remains unclear. Therefore, we first estimated the number and conductance of GABA_A_ receptors activated by optical stimulation of SOM-INs using non-stationary fluctuation analysis (Clements, 2003) before and after NMDA application (Figure 3A). This approach indicated that NMDAR activation produces an increase in GABA_A_ receptor number (before: 182 ± 49; after: 370 ± 87, n = 8, p=0.01) but not conductance (before: 41 ± 9 pS; after: 34 ± 6 pS, n = 8, p=0.32), suggesting that iLTP involves the addition of GABA_A_ receptors in the postsynaptic membrane. To test this hypothesis, we pharmacologically blocked SNARE-dependent insertion of receptors by including botulinum toxin type A (BoNT-A) in the patch pipette and found that this manipulation completely abolished iLTP (Figure 3B). Indeed, IPSCs were slightly reduced following NMDA application in cells loaded with BoNT-A (86 ± 4%, n = 8, p=0.01) while cells loaded with heat-inactivated BoNT-A (HI-BoNT) still exhibited iLTP (126 ± 3%, n = 7, p=0.02).

**Figure 3.**
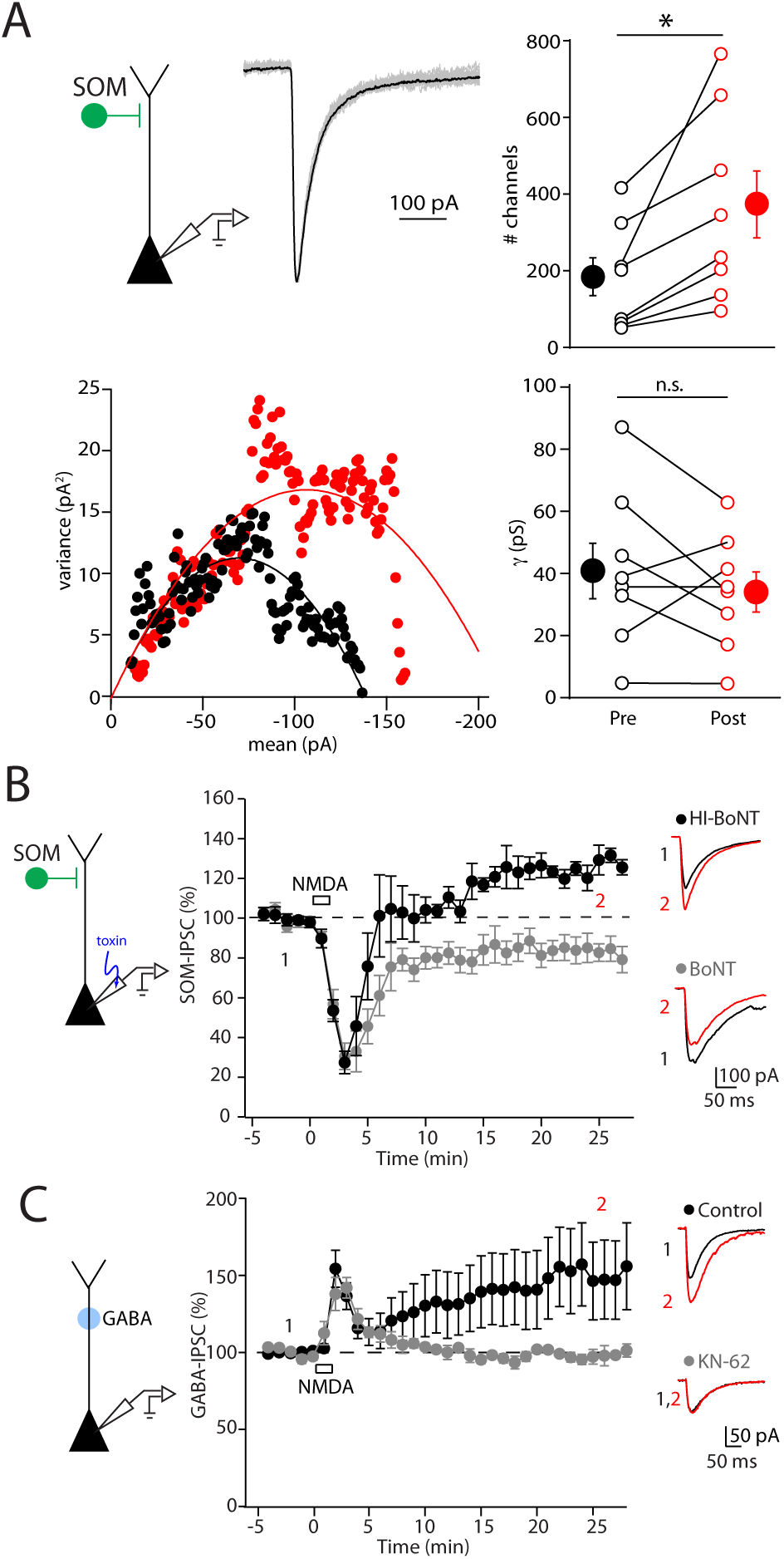
Dendritic iLTP is expressed postsynaptically. (A) Non-stationary fluctuation analysis of inhibitory responses evoked by photo-stimulation of SOM-INs in a representative experiment. Top, Example peak-scaled IPSC traces (gray) and average IPSC (black) during the baseline period. Bottom, Plot of the variance against the mean IPSC amplitude before (black) and after (red) NMDA application. Right, Summary plot of the estimated channel number (top) and conductance (bottom) before and after NMDA application. Asterisks denote p-value of <0.05, Paired T-test. (B) Time-course of inhibitory responses evoked by photostimulation of SOM-INs showing that botulinum type A (BoNT-A) abolishes iLTP (gray circles) while the heat-inactivated BoNT-A (black circles) does not. Right, Average IPSC traces obtained before (black) and after (red) NMDA exposure from single experiments are shown. (C) Time-course of IPSCs evoked by photolysis of RuBi-GABA at dendritic regions of L2/3 pyramidal cells. Under control conditions, IPSCs elicited by direct stimulation of postsynaptic receptors potentiate following NMDA application (black circles), and iLTP is blocked by KN-62 (gray circles). Right, Average IPSC traces obtained before (black) and after (red) NMDA exposure from single experiments are shown.

Finally, to further test the hypothesis that iLTP is expressed postsynaptically, we bypassed GABA release from presynaptic interneurons entirely and directly activated postsynaptic GABA_A_ receptors with photolysis of caged GABA targeting distal PN dendrites (Figure 3C). Consistent with postsynaptic iLTP expression, IPSCs evoked by GABA uncaging also increased following NMDA exposure (161 ± 29%, n = 10, p=0.0009), and this result was blocked by bath-application of KN-62 (99 ± 3%, n = 5, p=0.03 compared to control). In combination, these results strongly indicate that iLTP is both induced and expressed postsynaptically and is restricted to subsets of GABAergic synapses.

We next tested whether synaptic activity can also trigger NMDAR-dependent iLTP (Figure 4). To activate excitatory synapses targeting distal PN dendrites, theta-burst stimulation (TBS) was delivered through an extracellular stimulating pipette placed in layer 1 while depolarizing the postsynaptic cell to −20 mV. This initial pairing protocol resulted in a long-lasting depression of IPSCs evoked by optical stimulation of SOM-INs (Supplemental Figure 2). We hypothesized this result might be due to a decrease in presynaptic GABA release as a result of metabotropic glutamate receptor (mGluR)-induced endocannabinoid mobilization (Chiu et al., 2010). Indeed, when mGluRs were blocked with the antagonist MCPG, TBS pairing triggered iLTP of SOM-IPSCs (131 ± 8%, n =10, p=0.007, Figure 4A, Supplemental Figure 2B). Moreover, consistent with our data using NMDA application, TBS pairing did not alter IPSCs evoked by stimulation of PV-INs (97 ± 6%, n =9, p=0.54; Figure 4A). We confirmed that iLTP triggered by TBS pairing also requires NMDARs as it was abolished by bath-application of the NMDAR-antagonist CPP (Supplemental Figure 2B). If the depression observed with intact mGluR signaling is indeed expressed presynaptically, then bypassing GABA release with photo-uncaging should reveal iLTP following TBS pairing without the need for mGluR blockade. Consistent with this prediction, TBS pairing potentiated uncaging-evoked IPSCs in control recording solution (143 ± 15%, n = 6, p=0.03; Figure 4B). In summary, we find that synaptic activation of NMDARs is sufficient to selectively induce postsynaptic iLTP of SOM-IN-mediated IPSCs.

**Figure 4.**
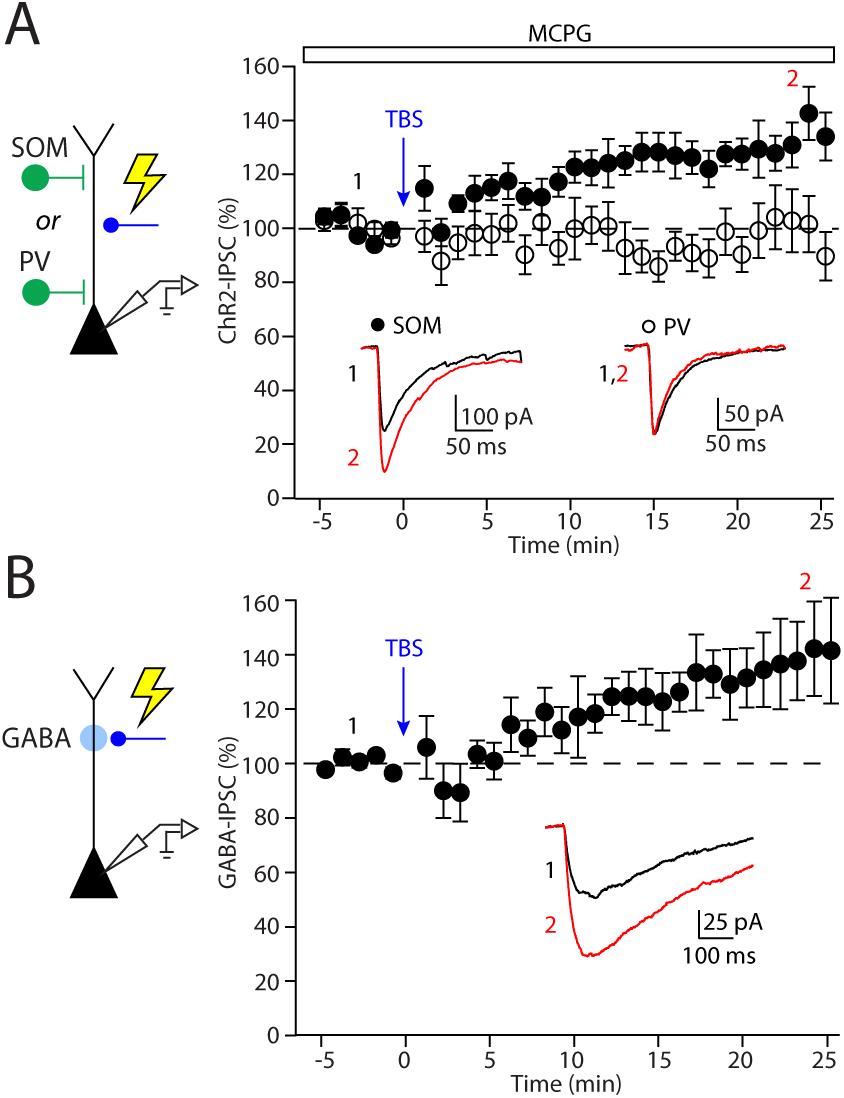
Excitatory synaptic stimulation induces iLTP of SOM-INs. (A) Time-course of inhibitory responses evoked by photostimulation of SOM- or PV-INs before (black) and after (red) theta-burst stimulation (TBS) in the presence of MCPG. Average IPSC traces are shown for representative experiments in the inset. (B) TBS triggers potentiation of IPSCs evoked by uncaging of RuBi-GABA in the absence of MCPG.

Our results suggest that the molecular constituency of synapses formed by SOM-INs may differ from those formed by other interneurons, resulting in differential sensitivity to plasticity induction. Previous studies have linked both β2 and β3 subunits of the GABA_A_ receptor to iLTP induction in the cerebellum and hippocampal cultures, respectively (He et al., 2015; Petrini et al., 2014). Therefore, we first asked whether functional expression of these subunits might distinguish synapses formed by SOM-versus PV-INs (Figure 5A). Bath application of etomidate, a β2/β3-selective positive allosteric modulator, slightly reduced the amplitude of IPSCs arising from both SOM- and PV-INs (SOM: −26 ± 6%, n = 7, p=0.02; PV: −17 ± 4%, n = 7, p=0.02). However, etomidate substantially slowed the decay of IPSCs evoked by optical stimulation of SOM-INs (baseline: 59 ± 9 ms; etomidate: 308 ± 102 ms, n = 7, p=0.02) but had no impact on the decay of PV-IN-evoked currents (baseline: 27 ± 1 ms; etomidate: 35 ± 4 ms, n = 7, p=0.16). This finding suggests that β2/β3 expression is enriched at synapses formed by SOM-INs versus PV-INs.

**Figure 5.**
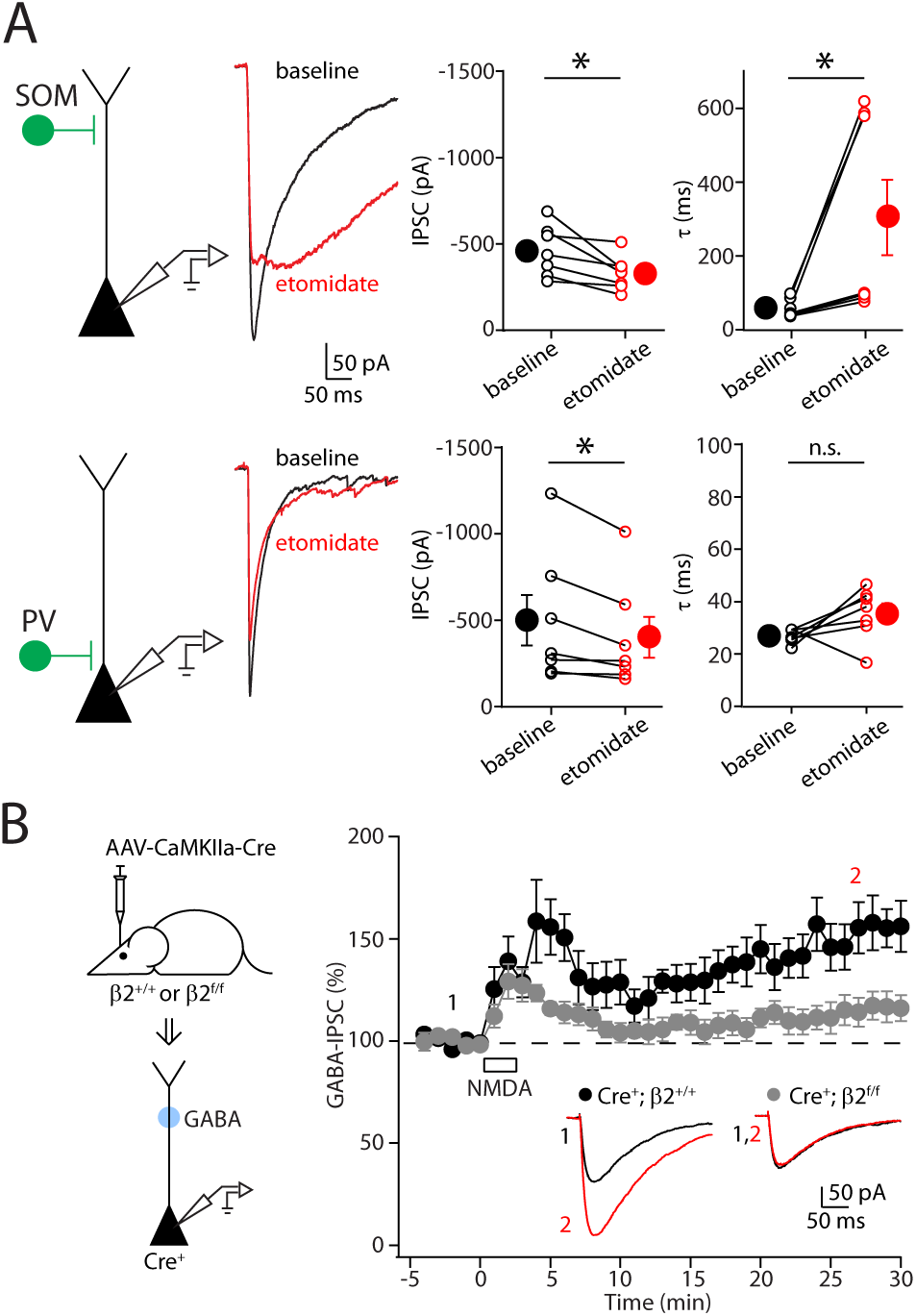
GABA_A_ receptor β2 subunits are enriched at synapses formed by SOM-INs and are required for iLTP. (A, top) Average IPSC traces obtained from optogenetic stimulation of SOM-INs (left) at baseline (black) and in etomidate (red) for a representative experiment. Summary plots of the effect of etomidate on average IPSC amplitude and decay kinetics are shown. (A, bottom) Similar experiments are shown for optogenetic stimulation of PV-INs. Asterisks denote p-value of <0.05, paired T-test. (B, left) Schematic of the experimental paradigm. Pyramidal cells expressing GFP-Cre are targeted for patching, and inhibitory responses are elicited by uncaging GABA in the dendritic regions. (B, right) Time-course of the effect of NMDA application on uncaging responses in cells obtained from wild-type (black) or β2-deleted (gray) cells. Average IPSC traces obtained before (black) and after (red) NMDA exposure from a single experiment are shown in the insets.

We then tested whether β2- or β3-containing GABA_A_ receptors are required for iLTP by using mice expressing floxed conditional alleles of either the β2 (Supplemental Figure 3A,B) or β3 (Ferguson et al., 2007) subunit of the GABA_A_ receptor. We virally introduced GFP-tagged Cre recombinase (AAV-CaMKIIα-GFP-Cre, Figure 5B) into the prefrontal cortex of conditional mice and prepared acute slices 6-7 weeks post-injection (Supplemental Figure 3C). Notably, genetic deletion of the β2 subunit eliminated iLTP of uncaging-evoked IPSCs following NMDA application (Cre^+^; β2^f/f^: 115 ± 7%, n=8, p=0.08, Figure 5B). In contrast, neither expression of GFP-Cre by itself (Cre^+^; β2^+/+^: 161 ± 13%, n=6, p=0.03, Figure 5B) nor deletion of the β3 subunit blocked potentiation of uncaging-evoked IPSCs (Cre^+^; β3^f/f^: 161 ± 21%, n=7, p=0.02, Supplemental Figure 3D).

The preceding results indicate that activation of NMDARs can acutely potentiate the strength of inhibition mediated by selective subsets of GABAergic interneurons. We next asked whether glutamatergic signaling also plays a role in regulating inhibitory potency *in vivo*. To address this possibility, we utilized a genetic strategy for sparsely eliminating NMDAR signaling in prefrontal neurons in the intact mouse. We used the same viral vector to express GFP-tagged Cre recombinase in mice harboring a floxed allele of the obligatory NR1 subunit of the NMDAR (Tsien et al., 1996). In slices prepared 6-7 weeks following injection, infected (GFP-positive) and non-transfected (GFP-negative) cells were intermixed. Whole cell recordings of PN pairs combined with local electrical stimulation confirmed that NMDARs were not functional in Cre-expressing cells (Figure 6A, B). In contrast, the amplitude of AMPAR-mediated excitation was not significantly altered by NR1 deletion (Cre^−^; NR1^f/f^: −78.9 ± 9.9 pA, Cre^+^; NR1^f/f^: −118.2 ± 29.6 pA, n = 10, p=0.43, Figure 6A, B).

**Figure 6.**
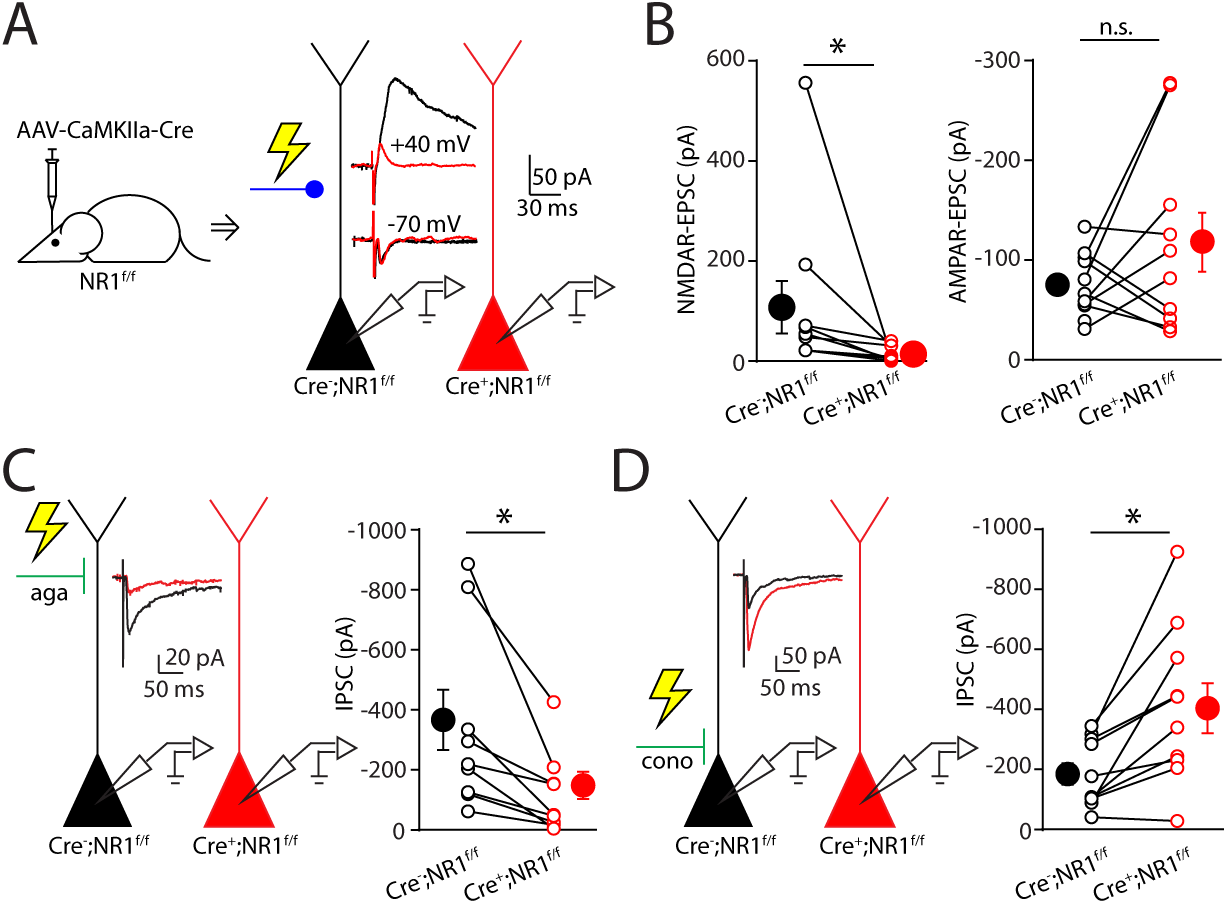
Conditional deletion of NMDARs differentially alters dendritic and perisomatic inhibition. (A) Schematic of whole-cell recordings in neighboring cells to compare evoked glutamatergic responses. AMPAR- (-70 mV) and NMDAR- (+40 mV) EPSCs in control (black) and NR1-lacking (red) cells are shown in the insets. (B) Summary plot of NMDAR- (left) and AMPAR-EPSC amplitude (right). (C, left) Schematic of dual recordings in neighboring cells to compare inhibitory responses evoked by an extracellular stimulating electrode in layer 1. Agatoxin (aga) was bath applied to block P/Q-type Ca2+ channels. Average IPSC traces are shown for a control (black) and a NR1-lacking (red) cell in a representative experiment. (C, right) Summary plot comparing amplitude of isolated IPSCs between wild-type and NR1-deleted cells. (D) Similar results obtained for IPSCs evoked by stimulating in layer 2/3 and in the presence of conotoxin (cono) to block N-type Ca2+ channels. Asterisks denote p-value of <0.05, Wilcoxon matched-pairs signed rank Test.

We then examined inhibition onto NR1-deleted cells. As Cre recombinase was utilized to remove NR1 expression, we could not adopt the same strategy of ChR2-mediated activation of IN subtypes. To compare inhibition putatively mediated by SOM-INs or PV-INs, we placed a stimulating electrode either in layer 1 or the cell body layer, respectively (Figure 6C, D). We further enhanced the selectivity of activation by including either the P/Q-type Ca2+ channel blocker agatoxin TK or the N-type channel blocker conotoxin GVIA in the bathing solution to block GABA release from specific INs (Kruglikov and Rudy, 2008). While PV-INs exclusively depend on P/Q-type Ca2+ channels for GABA release, SOM-INs utilize both channel types to mediate GABAergic transmission (Supplemental Figure 4A, B). Consistent with our preceding results, NR1 deletion led to a significant reduction in GABAergic inhibition putatively mediated by SOM-INs (Cre^−^; NR1^f/f^: −338.7 ± 100.4 pA, Cre^+^; NR1^f/f^: −120 ± 45.1 pA, n = 9, p=0.004, Figure 6C). Surprisingly, we found that loss of NMDAR signaling produced a significant enhancement of putative PV-IN-mediated inhibition (Cre^−^; NR1^f/f^: −187.1 ± 35.8 pA, Cre^+^; NR1^f/f^: −411.7 ± 83.2 pA, n = 9, p=0.004, Figure 6D). These results indicate that NMDARs control the strength of GABAergic inhibition *in vivo*, but the directionality of this influence differs across inhibitory synaptic subpopulations, potentially leading to a disruption of the balance between excitation and inhibition at the subcellular level.

## Discussion

The cellular mechanisms underlying the preservation of balance between synaptic excitation and inhibition across distinct GABAergic circuits remain poorly understood. Recent work has begun to focus on GABAergic synaptic plasticity as a key mediator of homeostatic control (Castillo et al., 2011; Kullmann et al., 2012). In this study, we found that activation of glutamatergic NMDARs by either exogenous agonists or endogenous glutamate is capable of potentiating GABAergic synapses in the neocortex. Notably, this form of plasticity is specific to inputs arising from SOM-INs due to the selective sensitivity of these dendritic inhibitory synapses to CaMKIIα activity. Indeed, loading single neurons with either calmodulin or constitutively active CaMKIIα enhanced responses to optical stimulation of SOM-INs but not PV-INs, suggesting that the postsynaptic composition of inhibitory connections is functionally and molecularly heterogeneous. Consistent with this hypothesis, pharmacological analysis suggested enrichment of the β2 subunit of the GABA_A_ receptor at inputs formed by SOM-INs, and genetic deletion of this subunit blocked induction of iLTP. Finally, we show that signaling through NMDARs is necessary *in vivo* for the maintenance of putative SOM-IN synapses. Our results highlight a novel mechanism for maintaining the balance of excitation and inhibition within neocortical dendrites.

Homeostatic regulation of GABAergic signaling in the neocortex has largely focused on perisomatic inhibition. Work *in vivo* showed that loss of visual stimulation resulted in a weakening of PV-IN synapses onto layer 4 PNs (Maffei et al., 2006). Similarly, the ratio of glutamatergic and GABAergic inputs to layer 2/3 PNs in visual cortex is highly conserved, despite large variations in the absolute magnitudes of each component (Xue et al., 2014). This balance was attributed to the regulation of synapses formed by PV-INs, as chronically altering PN spike output resulted in a corresponding change in perisomatic inhibition (Xue et al., 2014). In keeping with this observation, several studies demonstrated modulation of PV-IN inputs following alterations in pyramidal cell firing (Bartley et al., 2008; Holmgren and Zilberter, 2001). Indeed, postsynaptic spiking is sufficient to induce changes in inhibitory synaptic efficacy from fast-spiking, putative PV-expressing interneurons (Kurotani et al., 2008; Lourenco et al., 2014). Overall, these findings suggest a direct linkage of PN *output* and the strength of *perisomatic* inhibition.

In contrast to these studies, our results indicate a distinct relationship between excitation and inhibition, where glutamatergic *input* is coupled to the strength of *dendritic* GABAergic signaling. Supporting this idea, the coupling of NMDARs with GABAergic plasticity was previously shown in cultured hippocampal neurons (Marsden et al., 2007; Petrini et al., 2014), though input specificity was not addressed. Surprisingly, hippocampal iLTP was shown to require the β3 GABA_A_ subunit (Petrini et al., 2014), indicating that not all aspects of this phenomenon may be conserved. Nevertheless, it is intriguing to speculate that dendritic iLTP may be a general process across cortical areas. Given the dependence of SOM-IN iLTP on NMDAR-mediated Ca2+ influx, we predict that this form of plasticity will be highly localized in small dendritic regions, consistent with the compartmentalization of glutamatergic Ca2+ transients (Higley and Sabatini, 2008; Sabatini et al., 2002). We previously showed that inhibition mediated by SOM-INs could, in turn, influence excitatory transmission and Ca2+ signaling at the scale of single dendritic spines (Chiu et al., 2013), potentially driving long-term depression of glutamatergic inputs and regulating spine stability (Chen et al., 2015; Hayama et al., 2013). Thus, the homeostatic interaction of glutamatergic and GABAergic signaling may fine-tune excitatory synaptic integration at the level of individual synapses.

The critical role of inhibitory plasticity *in vivo* is also supported by recent work showing that GABAergic synapses formed in the dendrites of L2/3 PNs of visual cortex are highly dynamic both spontaneously and in response to altered sensory experience (Chen et al., 2012; Kannan et al., 2016; Villa et al., 2016). Notably, GABAergic inputs to distal dendrites exhibit greater turnover than more proximal contacts, with synapses on dendritic spines among the most labile (Chen et al., 2012). This observation is consistent with our earlier findings that SOM-INs make a subset of their inputs directly onto spine heads (Chiu et al., 2013). Given these results and our present findings, it would be interesting to examine the role of NMDARs in visual experience-dependent reorganization of cortical GABAergic circuits.

The mechanisms underlying molecular heterogeneity of GABAergic synapses are unclear and likely involve the differential trafficking of receptor subunits and accessory molecules to distinct pools of synapses across the somatodendritic arbor. In contrast to glutamatergic synapses, the structural organization of GABAergic inputs is not well characterized. Previous work has suggested the possibility that inhibitory scaffolding molecules may vary across synaptic subpopulations. In the neocortex, the cell adhesion molecule neuroligin-2 was reported to be necessary for synapses formed by PV-INs but not SOM-INs (Gibson et al., 2009) In the cerebellum, the scaffolding molecule gephyrin was suggested to be critical for dendritic but not perisomatic GABAergic inputs to Purkinje cells (Viltono et al., 2008). Recent studies have begun to reveal additional molecules involved in the structure and function of inhibitory synapses (Uezu et al., 2016; Yamasaki et al., 2017), and future investigation will be necessary to determine their selective roles in different cellular compartments.

Previous models of synaptic homeostasis often rely on a straightforward “balance” of overall excitation and inhibition that may be oversimplified. As we have shown, dysregulation of NMDAR signaling results in opposite alterations in putative PV- and SOM-IN-mediated inhibition, possibly due to the uncoupling of glutamatergic synaptic signaling and somatic spiking. This process results in a redistribution of inhibition along the somatodendritic axis. Many studies have suggested that the functional roles of inhibition mediated by different IN populations are highly distinct (Atallah et al., 2012; Lee et al., 2012; Wilson et al., 2012). Thus, although the total amount of inhibition may remain “balanced”, the functional consequences for cellular and circuit activity may be considerable.

In conclusion, we present evidence for a novel synapse-specific mechanism for linking excitatory signaling to the potency of dendritic GABAergic inhibition. We expect that future studies into the cellular mechanisms governing such specificity will yield rich rewards into understanding both basic synaptic development and maintenance as well as circuit organization and function.

## Methods

### Slice Preparation

All animal handling was performed according to the Yale Institutional Animal Care and Use Committee and federal guidelines. Optogenetic ChR2 experiments were performed in acute prefrontal cortical (PFC) slices taken from specific interneuron Cre-driver lines (SOM-Cre, PV-Cre or VIP-Cre mice)(Taniguchi et al., 2011) at postnatal day (P) 30-50 expressing ChR2 in targeted IN populations. GABA uncaging experiments were conducted using cortical slices from wild-type C57/Bl6 mice (P30-50) or transgenic mice harboring floxed alleles of GluN1, Gabrb2, or Gabrb3 (P55-70). Under isoflurane anesthesia, mice were decapitated and coronal slices (300 μm thick) were cut in ice-cold external solution containing (in mM): 100 choline chloride, 25 NaHCO_3_, 1.25 NaH_2_PO_4_, 2.5 KCl, 7 MgCl_2_, 0.5 CaCl_2_, 15 glucose, 11.6 sodium ascorbate and 3.1 sodium pyruvate, bubbled with 95% O2 and 5% CO_2_. Slices containing the prelimbic-infralimbic regions of the PFC were then transferred to artificial cerebrospinal fluid (ACSF) containing (in mM): 127 NaCl, 25 NaHCO3, 1.25 NaH2PO4, 2.5 KCl, 1 MgCl2, 2 CaCl2 and 15 glucose, bubbled with 95% O2 and 5% CO2. After an incubation period of 30 min at 34 °C, the slices were maintained at 22–24 °C for at least 20 min before use.

### Electrophysiology

Experiments were conducted at room temperature (22–24 °C) in a submersion-type recording chamber. Whole-cell voltage-clamp recordings were obtained from layer 2/3 pyramidal cells located 200-300 μm from the pial surface and identified with video infrared differential interference contrast. To obtain measurable GABA_A_R responses at a membrane holding potential of −70 mV, glass electrodes (3.0-3.2 MΩ) were filled with a high chloride internal solution containing (in mM): 100 CsCl, 30 CsGluconate, 10 HEPES, 4 MgCl_2_, 4 Na_2_ATP, 0.4 NaGTP and 10 sodium creatine phosphate, adjusted to pH 7.3 with CsOH. In GABA uncaging experiments, red-fluorescent Alexa Fluor-594 (10 μM)(Invitrogen) was included in the pipette solution to visualize cell morphology. For all recordings, series resistance was 15-25 MΩ and uncompensated. Recordings were discarded if series resistance changed >15% during the experiment. Electrophysiological recordings were made using a Multiclamp 700B amplifier (Molecular Devices), filtered at 4 kHz, and digitized at 10 kHz using acquisition software written in Matlab (Mathworks)(Pologruto et al., 2003).

### Synaptic stimulation and GABA uncaging

To photoactivate specific INs, SOM-Cre, PV-Cre or VIP-Cre mice were injected at P14-20 into the PFC with recombinant AAV driving Cre-dependent expression of a ChR2-EYFP fusion protein under the Ef1α promoter (AAV-DIO-Ef1α-ChR2-EYFP)(University of North Carolina Vector Core). Mice were sacrificed 14-21 days post-injection for slice preparation as described above. To activate ChR2-positive fibers, we filled the back aperture of the microscope objective (60x, 1.0 NA, Olympus) with blue light from a fiber-coupled 473 nm laser (Optoengine LLC), yielding a ^~^15-20 μm diameter disc of light at the focal plane. A brief (0.5-3 ms) pulse of light (3-5 mW at the sample) reliably stimulated ChR2-expressing INs and evoked IPSCs in pyramidal neurons. To photorelease GABA, 11 μM Rubi-GABA (Chiu et al., 2013; Rial Verde et al., 2008) was included in the bathing ACSF and the microscope objective was centered over the apical dendritic arbor of the recorded neuron. Light pulses were delivered as with optogenetic stimulation and reliably evoked IPSCs. For experiments involving local electrical stimulation, a glass theta stimulating electrode was placed in layer 1 or 2/3 to evoke IPSCs in distal or perisomatic regions, respectively. For theta burst stimulation (TBS) of excitatory afferents, a monopolar stimulating electrode was placed in either layer 1 or layer 2/3, depending on the target site (dendritic versus perisomatic) of inhibitory plasticity. TBS consisted of 5 bursts (4 pulses at 100 Hz) delivered at 5 Hz, repeated 10 times every 5 s. Patched cells were depolarized to −20 mV during bursting.

### Conditional deletion of targeted receptor subunits

To remove functional NMDARs, mice harboring a floxed allele of GluN1 (Tsien et al., 1996) (P14-20) were injected into the PFC with AAV driving expression of a Cre-GFP fusion protein under the CaMKIIαpromoter (AAV-CaMKIIα-Cre-GFP)(University of North Carolina Vector Core). Virus was diluted 1:10 and injected at a volume of 1 μl to obtain sparse infection. Mice were sacrificed 6-7 weeks post-injection for slice preparation as described above. A similar approach was used to delete the GABA_A_ receptor β2 (see below) or β3 (Ferguson et al., 2007) subunit.

### Generation of conditional β2 knockout mice

A BAC clone containing the GABA_A_R β2 gene (Gabrb2) from C57BL/6 mice genomic DNA was purchased from BACPAC Resources Center (Oakland, CA USA). We combined MultiSite Gateway cloning technology (Invitrogen, Carlsbad, CA USA) and Red/ET-mediated homologous recombination (Gene Bridges GmbH, Heidelbelg, Germany) for targeting vector construction. The targeting vector was linearized, electroporated into the embryonic stem (ES) cell line RENKA derived from the C57BL/6N strain (Mishina and Sakimura, 2007), and selected by G418. Recombinant clones were identified by Southern blot analysis using the *Gabrb2* 5’ probe on Spe I-digested genomic DNA, and the *Gabrb2* 3′ probe on BamH I-digested genomic DNA. Targeted clones were injected into eight-cell stage embryos of a CD-1 mouse strain. The embryos were cultured to blastocysts and transferred to pseudopregnant CD-1 mice. Resulting male chimeric mice were crossed with female C57BL/6N mice. After Cre-loxP recombination, the elimination of exon 4 results in a frame-shift mutation in the gene encoding GABA_A_R β2.

### Data analysis

Off-line analysis of electrophysiological recordings was performed using custom routines written in IgorPro (Wavemetrics). IPSC amplitudes were calculated by finding the peak of the current traces and averaging the values within a 1 ms window. Potentiation of GABAergic responses was assessed by comparing the average IPSC amplitude in the first 5 minutes prior to NMDA application to the average IPSC amplitude 20-25 minutes after NMDA washout for each experiment, using paired Student’s t-tests at a significance level of p<0.05. To assess the effect of pharmacological blockade on iLTP, we performed Mann-Whitney tests comparing drug versus control experiments. For recordings comparing pairs of neighboring GluN1-positive and -negative cells, a Wilcoxon matched-pairs signed rank test was performed to assess significance at p<0.05 due to non-normally distributed data.

### Pharmacology

For all optogenetic experiments, the ACSF included 10 μM NBQX to block AMPA receptors. In a subset of experiments (see text), the ACSF also included (in μM): 6 ifenprodil, 3 CGP-55845, 3 nimodipine, 5 KN-62, 20 cyclosporine A, 100 (S)-MCPG, 50 CPP, 0.5 etomidate, 0.2 ω-agatoxin TK or 1 ω- conotoxin GVIA. In cell loading experiments, the drug concentrations (in μM unless indicated otherwise) are as follows: 500 MK-801, 10 mM BAPTA, 10 AIP, 40/10 Ca2+/calmodulin or 200 ng/ml BoNT-A. For loading constitutively active CaMKII*, the compound was synthesized as previously described (Tavalin and Colbran, 2017) and added to the internal solution. All compounds other than CaMKIIα were purchased from Tocris except for conotoxin (Peptides International), agatoxin (Peptides International), and calmodulin (Sigma-Aldrich).

## Author Contributions

CYC and MJH designed the experiments and wrote the manuscript. CYC performed all experiments. JSM, ST, MY, and RN designed and generated the conditional β2 subunit mouse. SJT designed and synthesized constitutively active CaMKIIα.

## Acknowledgements

The authors wish to thank Dr. Jessica Cardin for helpful comments during the preparation of this manuscript. This work was funded by the NIH (R01 MH099045 to MJH) and the March of Dimes Basil O’Connor Award to MJH.

---

**Supplemental Figure 1.**
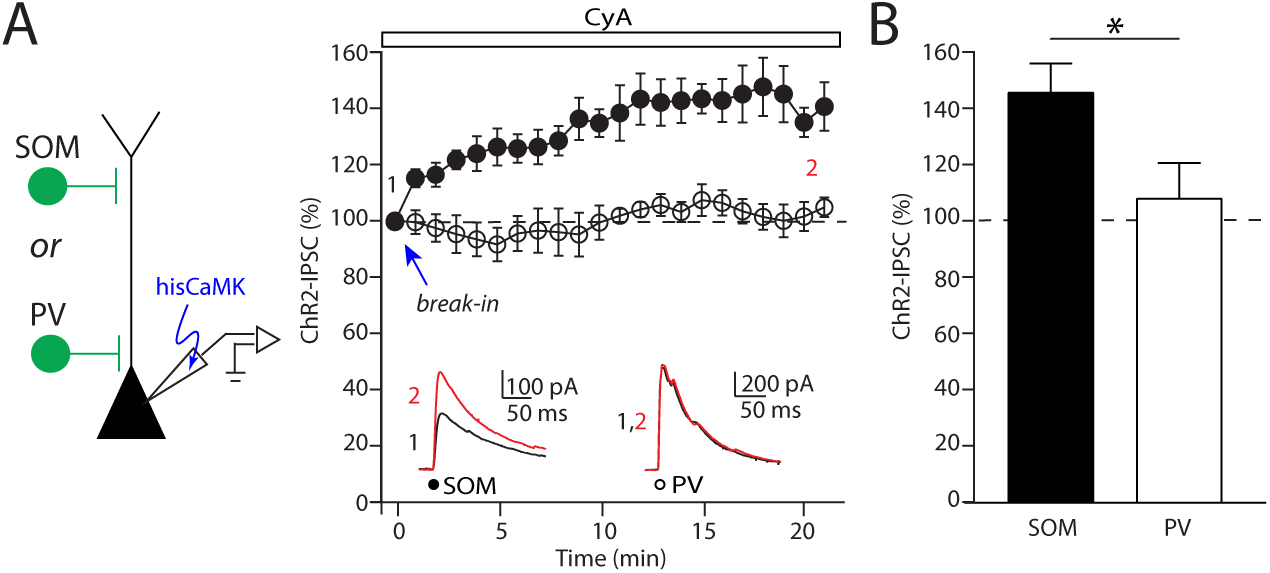
Loading activated CaMKIIα specifically potentiates GABAergic inhibition from SOM-INs. (A) Time-course of inhibitory postsynaptic currents evoked by photo-activation of SOM or PVINs immediately after whole-cell break-in using an internal patch solution that contains activated CaMKIIα. Calcineurin was blocked with bath application of cyclosporin A (CyA) throughout the experiment. Average IPSC traces obtained in the first minute (black) and after twenty minutes (red) of whole cell patch recording from a single experiment are shown in the insets. (B) Summary plot of the effect of loading activated CaMKIIα on the amplitude of inhibitory responses elicited by optogenetic activation of SOM- and PV-INs. Asterisks denote p-value of <0.05, Mann-Whitney Test.

**Supplemental Figure 2.**
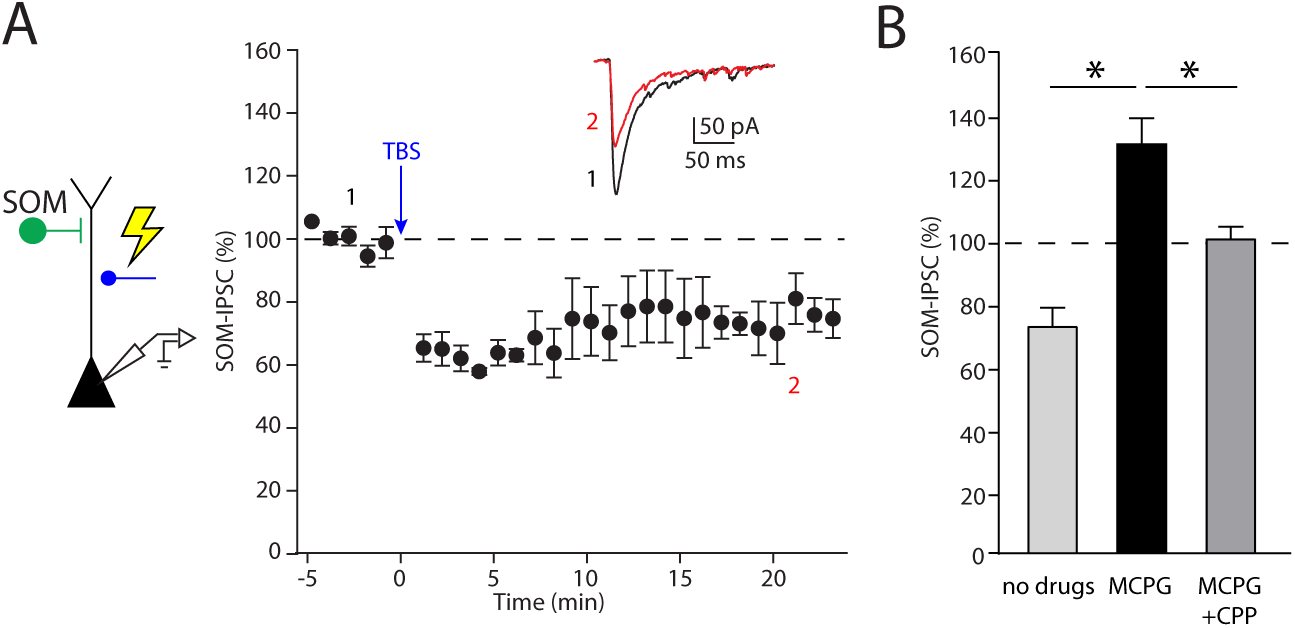
Synaptic stimulation of excitatory afferents induces iLTD of SOM-INs with intact mGluR activity. (A) Time-course of inhibitory responses evoked by photostimulation of SOM-INs before (black) and after (red) theta-burst stimulation (TBS) in control ACSF. Average IPSC traces are shown for a representative experiment in the inset. (B) Summary plot of the effect of TBS on inhibition from SOM-INs in different recording solutions. Asterisks denote p-value of <0.05, Mann-Whitney Test.

**Supplemental Figure 3.**
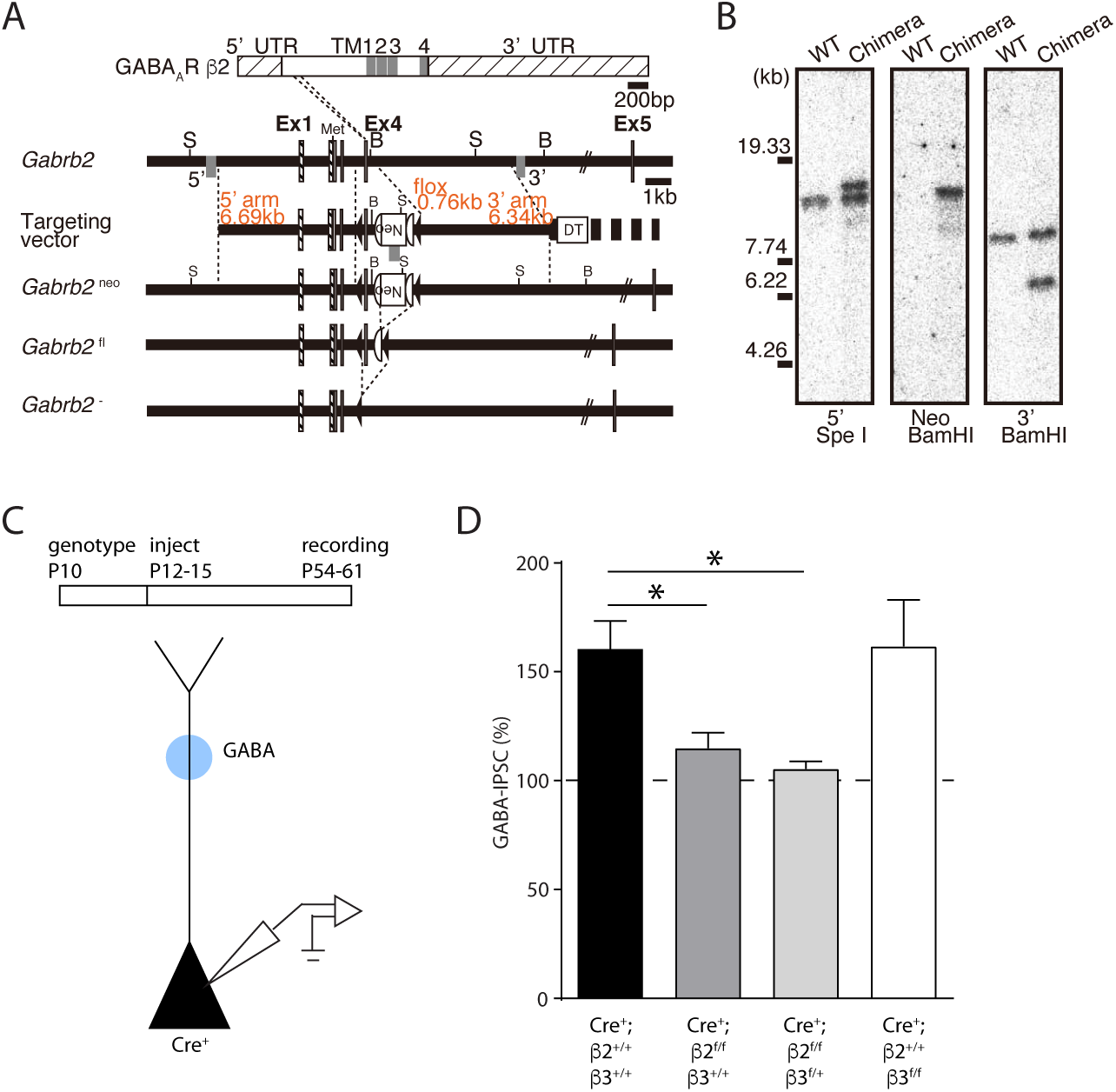
Differential role of distinct β subunits in iLTP. (A,B) Schematic illustrating design of conditional β2 mouse. (C) Schematic of the experimental paradigm showing the timing of genotype determination, viral injection and electrophysiological assessment. Pyramidal cells expressing GFP-Cre are targeted for patching, and inhibitory responses are elicited by uncaging GABA in the dendritic regions. (D) Summary plot of the effect of NMDA application on uncaging responses in cells obtained from mice in which expression of β2 (dark gray bar) or β3 (white bar) or both subunits (light gray bar) are altered. Asterisks denote p-value of <0.05, Mann-Whitney Test.

**Supplemental Figure 4.**
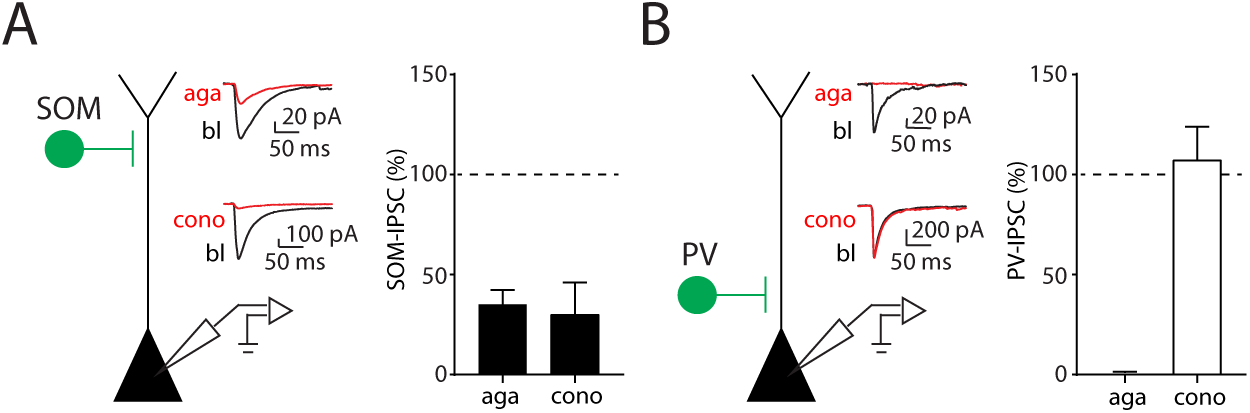
Voltage-gated Ca2+ channel blockers differentially modulate inhibition mediated by SOM- or PV-INs. (A) Schematic of whole-cell recordings in L2/3 pyramidal cells to assess the impact on optogenetically activated inhibitory responses from SOM-INs. Average IPSC traces from single experiments are shown in the inset for the effect of agatoxin (top) and conotoxin (bottom). Summary plots of the percent of IPSC amplitude in the respective toxins (red traces) relative to baseline (black traces) are presented to the right. Aga: 35.0 ± 7.3%, n=3, p=0.01; Cono: 30.3 ± 15.7%, n=4, p=0.02. (B) Similar experiments performed for PV-INs. Aga: 1.4 ± 0.0007%, n=4, p<0.0001; Cono: 107.5 ± 16.5%, n=3, p=0.70.

